# A genome-wide CRISPR screen supported by human genetics identifies the *TNRC18* gene locus as a novel regulator of inflammatory signaling

**DOI:** 10.1101/2023.10.04.560902

**Authors:** Fedik Rahimov, Sujana Ghosh, Sakina Petiwala, Mary Schmidt, Eugene Nyamugenda, Jason Tam, Daniel Verduzco, Sanjana Singh, Victor Avram, Apexa Modi, Celso A. Espinoza, Charles Lu, Jing Wang, Ashleigh Keller, Michael Macoritto, Naim Al Mahi, Tifani Anton, Namjin Chung, Michael J. Flister, Kanstantsin V. Katlinski, Amlan Biswas, Anneke I. den Hollander, Jeffrey F. Waring, Joshua D. Stender

## Abstract

Interleukin-1β (IL-1β) is dysregulated in many chronic inflammatory diseases, yet the genetic factors influencing IL-1β production and signaling remain largely unknown. Myeloid-derived cells are the primary producers of IL-1β, prompting a genome-wide CRISPR knockout screen in the human myeloid-derived U937 cell model, treated with lipopolysaccharide (LPS) to mimic inflammatory conditions, and sorted for high and low intracellular IL-1β levels. A total of 295 genes were identified as regulators of IL-1β production, including known mediators, such as TLR4, JAK-STAT, IL-10 receptor, and the Cullin ring finger ligase complex. Notably, 57 out of the 295 genes overlapped with loci associated with human inflammatory diseases, including the *TNRC18* gene on chromosome 7p22.1 associated with multiple diseases in the Finnish population. U937 cells engineered with the homozygous rs748670681 risk allele associated with inflammatory bowel disease, demonstrated decreased levels of mRNA for *TNRC18* and an adjacent gene *WIPI2*, reduction in LPS-dependent gene activation and cytokine production, but elevation of interferon-responsive gene programs. Transcriptomic profiles for individual knockouts of *TNRC18* and *WIPI2* attributed the loss of LPS-dependent signaling primarily to *TNRC18* while the exacerbation of interferon signaling is a hallmark of loss of *WIPI2*. Collectively, these findings delineate the global regulatory mechanisms of IL-1β production and provide molecular insights to the role of the rs748670681 variant as a pleiotropic risk factor for inflammatory diseases.

## INTRODUCTION

Precise regulatory control of the innate and adaptive immune responses is essential for proper tissue homeostasis and the prevention of inflammation-driven diseases. Cytokines play a critical role in the maintenance of this homeostasis. Sustained levels of inflammatory cytokines such as TNF-α, IL-6, and IL-1β contribute to the development of chronic diseases including type II diabetes^1^, atherosclerosis^2^, cancer^3^, neurodegeneration^4^, and autoimmunity^5^. These cytokines are typically produced by hyperactivated effector immune cells such as macrophages, natural killer cells, B cells, and dendritic cells. Establishing and maintaining immune cells throughout hematopoietic differentiation requires coordinated activities of pioneering and signal-dependent transcription factors^6^. For macrophages, PU.1 is an essential factor that binds to specific DNA sequences within regulatory regions, reorganizes the nucleosome structure, and interacts with members of the AP1 and C/EBP family of transcription factors to establish a permissive epigenetic landscape, including high levels of H3K27ac and H3K4me1/2/3^7,8^. In response to pathogen-associated molecular patterns recognized by pattern recognition receptors (PRRs) or damage-associated molecular patterns (DAMPs), signal-dependent transcription factors including members of the NFκB, STAT, and IRF families bind to active regulatory regions established by PU.1, recruit the RNA polymerase complex, and initiate rapid and robust activation of proinflammatory gene programs^9^.

One example of this regulation is the proinflammatory cytokine IL-1β, essential for maintaining immune homeostasis, whose dysregulation is a hallmark of chronic inflammation. As such, PU.1 constitutively binds to the *IL1B* promoter and is a pioneering factor to allow LPS-dependent recruitment of the NFκB family to promote transcriptional activation^10^. Pro-IL-1β, the precursor for IL-1β, is encoded by the *IL1B* gene located in a cluster of IL1 genes on chromosome 2. Upon activation of PPRs, the mRNA for IL-1β is rapidly increased in myeloid cells. Pro-IL-1β is synthesized intracellularly as a relatively inactive 31 kDa protein until processed into mature IL-1β by caspase-dependent inflammasomes and secreted from the cell through a variety of mechanisms, including exocytosis of secretory lysosomes, shedding of plasma membranes, and pyroptosis^11^. IL-1β interacts with the IL1 receptors on target cells to activate the NFκB, JNK, and p38 signaling pathways to induce downstream inflammatory genes^12^.

Over the past two decades, hundreds of genome-wide association studies (GWAS) have established the link between genetic variation and human immune-mediated diseases^13,14^. Interestingly, shared genetic associations have been observed for multiple immune diseases^15–17^, supporting the idea of hubs and hot spots driving the origin of multiple immune disorders^18^. Recent studies have demonstrated that therapeutic targets with genetic linkage to diseases have increased success rates in clinical trials and for reaching FDA approval^19–22^.

Gene products with roles in multiple diseases are particularly attractive therapeutic targets as they can be repurposed for numerous indications. For example, TNF-α, JAK1/2, IL-12B, TYK2, and IL-4R have been successfully targeted and gained regulatory approvals for inflammatory diseases such as psoriasis^23,24^, inflammatory bowel disease^25–27^, and atopic dermatitis^28,29^. Not all therapeutic agents equally demonstrate desirable efficacies for patients, necessitating the discovery of effective therapeutic agents for refractory patient populations. Therefore, focusing on targets with substantial phenotypic activities, appropriate cellular expression levels, and genetic associations to human diseases is cost-effective. IL-1 blockade therapies using anakinra have been successful in the clinic for treating patients with psoriatic arthritis, ankylosing spondylitis, and rheumatoid arthritis^30^. However they have been unsuccessful in treating systemic lupus erythematosus and Sjögren’s syndrome^31^. Upstream factors involved in IL-1β regulation might provide better therapeutic targets for additional autoimmune indications.

In this study we aimed to utilize unbiased genome-wide CRISPR-Cas9 mediated loss-of-function screening to identify regulators of IL-1β production and processing in LPS-treated myeloid cells. We identify known and novel regulators of IL-1β intracellular expression, with 57 showing genetic association with at least one immune indication. Furthermore, we provide evidence that one of the identified IL-1β regulators, Trinucleotide Repeat Containing 18 (TNRC18), harbors a functional intronic single nucleotide polymorphism (SNP) associated with multiple immune-mediated diseases in the Finnish population^32^.

## RESULTS

### Genome-wide CRISPR knockout screen identifies modulators of intracellular IL-1β expression

To assess the robustness of Toll-like receptor 4 (TLR4; the receptor for LPS)-dependent activation and secretion of IL-1β in the human monocytic cell line U937, we treated phorbol-12-myristate-13-acetate (PMA)-differentiated U937 cells with increasing doses of LPS and measured secreted levels of IL-1β. Secretion of IL-1β correlated with the LPS dose and reached maximum concentration at around 500 pg/ml. However, the addition of Nigericin, a potent activator of the NLRP3 inflammasome which promotes the processing and release of IL-1β, robustly enhanced IL-1β secretion to 12,000 pg/ml starting at a dose of 1 mg/ml LPS (Fig. 1a). In addition, we observed a 100-fold increase in intracellular levels of IL-1β after 24 hours of LPS stimulation (Fig. 1b). Therefore, exposure of PMA-treated U937 cells to LPS leads to robust production of IL-1β protein expression, which remains primarily intracellular and is secreted upon activation of the inflammasome.

**Figure 1:**
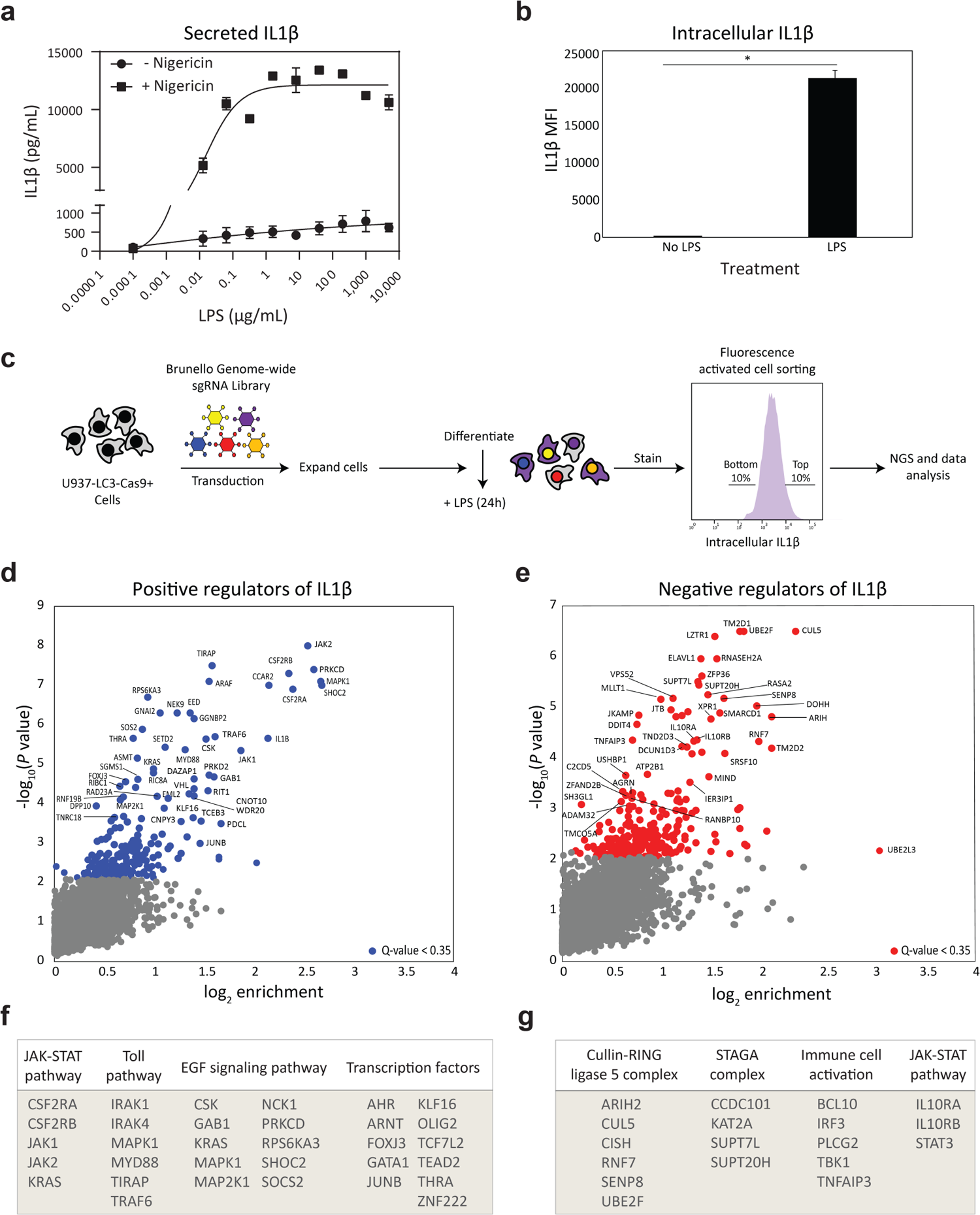
Pooled CRISPR knockout screen of IL-1β regulators in human monocytic cells U937. **a**) Treatment of PMA-differentiated U937-LC3-Cas9+ cells with increasing doses of LPS induces expression and secretion of IL-1β. Addition of Nigericin augments the secretion of IL-1β. **b**) Intracellular IL-1β level upon induction with LPS compared to vehicle treatment. **c**) Overview of the genome-wide CRISPR screening strategy using the Brunello sgRNA library. **d-e**) Genes that increase (**d**) or decrease (**e**) IL-1β levels when knocked out in U937 cells. For each gene the x axis shows the enrichment of sgRNAs in log_2_ (Q) values, and the y axis shows statistical significance of enrichment as -log_10_ (*P* value). **f-g**) Pathways and classes of known and novel genes that modulate IL-1β levels identified in IL-1β^low^ (**f**) and IL-1β^high^ (**g**) populations. LPS, lipopolysaccharide; MFI, median fluorescence identity; NGS, next-generation sequencing. Statistical significance was determined using two-tailed, unpaired Student’s *t* test (\**P* < 0.05).

To conduct genetic screens, U937 cells expressing constitutive Cas9 and a LC3 autophagy reporter (U937-LC3-Cas9+)^33^ were transduced with the pooled Brunello CRISPR library containing 76,441 single guide RNAs (sgRNAs) systematically targeting every protein-coding gene (n = 19,114) in the genome, each with four independent sgRNAs, and 1,000 non-targeting controls^34^. Transduced cells were differentiated with PMA and treated with LPS for 24 hours. Cell populations with the lowest 10% or highest 10% levels of intracellular IL-1β were collected using fluorescence-activated cell sorting (FACS), and sgRNA counts were compared in each population to pre-sorted control population with next-generation sequencing (Fig. 1c). We performed this experiment in triplicate and considered genes to be significantly enriched if they had a Q-value < 0.35 in either high or low sorted populations when compared to presort samples in 2 or 3 replicates. This analysis identified 128 genes (Q-value < 0.35) (Fig. 1d, Supplementary Table 1), whose CRISPR-mediated knockout led to loss of IL-1β production, and 167 genes (Q-value < 0.35) that led to increased intracellular levels of IL-1β (Fig. 1e, Supplementary Table 2).

As expected, members of the TLR4 pathway including IRAK1, IRAK4, MYD88 and TRAF6 were enriched in the low IL-1β population (Fig. 1f). In addition, members of the JAK-STAT pathway (CSF2RA/B, JAK1, JAK2), EGF pathway (e.g., KRAS, MAPK1, MAP2K1, SHOC2), and many transcription factors (e.g., JUNB, AHR, GATA1) were essential for LPS-dependent activation of IL-1β. Conversely, knockout of members of the Cul5 ubiquitin ligase (ARIH2, CUL5, RNF7, SENP8, UBE2F3) and the STAGA (101, KAT2A, SUPT7L, SUPT20H) complexes and genes involved in immune cell activation (BCL10, IRF3, TNFAIP3, TBK1) lead to elevated intracellular IL-1β (Fig. 1g). Collectively, these unbiased genome-wide screening results confirm known pathways of IL-1β production and identified several dozen novel regulators of IL-1β.

### Arrayed screen validates regulators of IL-1β production and secretion

Our pooled CRISPR screen was designed to capture genes that modulate intracellular IL-1β protein levels measured through flow cytometry. However, the identified genes may modulate IL-1β levels at various stages of regulation including production, processing, or secretion. To confirm our genome-wide pooled screening results and provide additional mechanistic support, we selected 18 genes with lowest and 18 genes with the highest levels of intracellular IL-1β for a multiplexed arrayed CRISPR screen to evaluate the effect of gene knockout on two phenotypic readouts: 1) intracellular IL-1β production through flow cytometry and 2) production and secretion of IL-1β using Meso Scale Discovery (MSD) immunoassays in supernatants. We confirmed all 18 screening hits identified as positive regulators of intracellular IL-1β levels, including expected regulators such as IL-1β, TRAF6, and MYD88, in addition to novel regulators such as CCDC134, RIT1, and CSK (Fig. 2a). CRISPR knockout of these genes also reduced the secretion of IL-1β in the presence of LPS or LPS and Nigericin, supporting a role for these genes in IL-1β mRNA expression or protein production (Fig. 2c). Conversely, knockout of 12 of 18 genes that had the highest levels of IL-1β in the pooled CRISPR screen, showed enhanced intracellular IL-1β levels in the arrayed screen; however, the majority (13 of 18) of these screening hits reduced IL-1β secretion in supernatants compared to control samples. For example, knockout of several members of the CRL5 complex (*CUL5*, *RNF7*, *ARIH2*) led to robustly increased intracellular IL-1β (Fig. 2b) but reduces the secretion of IL-1β even in the presence of Nigericin (Fig. 2d). Therefore, knockout of these genes leads to elevated intracellular IL-1β levels that fail to be secreted from the cell even upon activation of the inflammasome.

**Figure 2:**
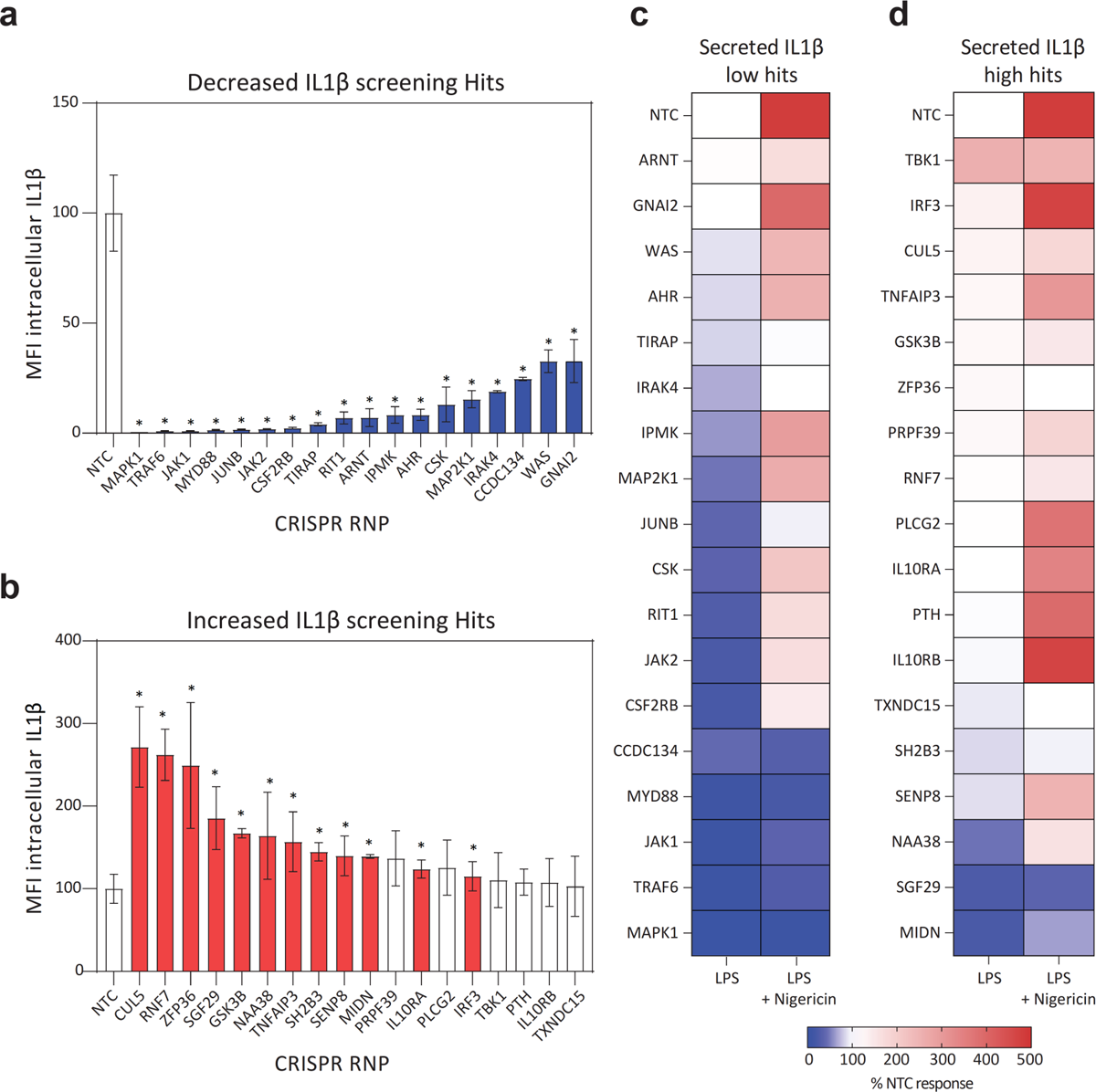
Arrayed CRISPR validation of IL-1β regulators in human monocytic cells U937. **a-b)** Intracellular IL-1β production measured by flow cytometry following LPS treatment of U937-LC3-Cas9+ cells that were transduced with sgRNAs targeting genes identified in genome-wide CRISPR screens in either (**a**) low or (**b**) high IL-1β populations. **c-d**) Normalized levels of secreted IL-1β collected from supernatants of LPS or LPS + Nigericin treated U937 cells following Cas9-RNP nucleofections targeting of CRISPR screening hits for low and high IL-1β sorted populations. MFI, median fluorescence identity; RNP, ribonucleoprotein; NTC, non-template control; LPS, lipopolysaccharide. Statistical significance was determined using two-tailed, unpaired Student’s *t* test (\**P* < 0.05) comparing NTC to CRISPR targeted cells.

### Genetic associations of CRISPR screening hits with immune-mediated diseases

Despite extensive efforts to fine-map GWAS loci, identifying the underlying target genes remains challenging. Genes that regulate the expression and/or secretion of an essential cytokine, such as IL-1β, can provide supporting evidence for causal genes at GWAS loci implicated in immune-mediated diseases. The Open Targets “locus-to-gene” (L2G) model ranks genes at each GWAS locus using genetics and a range of epigenetic features^35^. To assess the immune disease association of genes identified in our CRISPR screen, we queried the Open Targets Genetics database for supporting evidence for these genes at GWAS risk loci. We selected 93 published GWAS and studies conducted on immune-mediated diseases in FinnGen and UK Biobank in which loci containing top screening hits showed genome-wide significant associations. Harmonization of identical (e.g., atopic dermatitis and eczema combined as atopic dermatitis) or related phenotype (e.g., ACPA-positive rheumatoid arthritis [RA] and [RA] combined as [RA]) definitions across studies resulted in 28 unique or jointly evaluated phenotypes (Supplementary Table 3). Among the 295 screening hits 57 genes (19%) reside in loci associated with at least one immune disease and linked to GWAS signals with an average L2G score of 0.1 and above (Fig. 3a, Supplementary Table 3). Twenty-eight genes were linked to GWAS signals with a hard threshold^35^ of L2G ζ 0.5.

**Figure 3:**
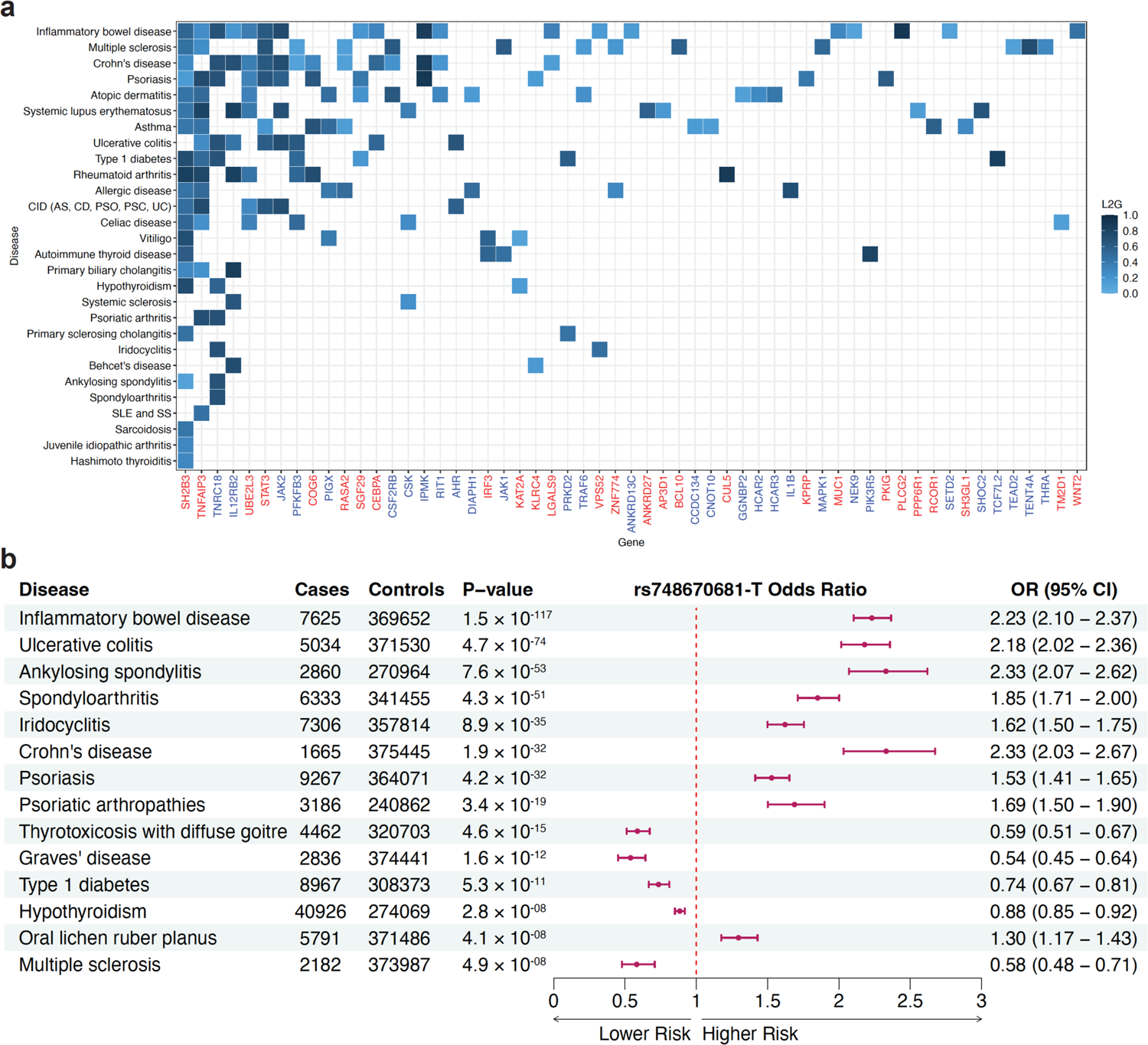
Genetic association of CRISPR screening hits with immune-mediated diseases. **a**) Top locus-to-gene (L2G) scores from Open Targets Genetics for 57 gene-disease pairs. Fifty-seven genes identified in the CRISPR screen were linked to various immune-mediated diseases in GWAS with varying levels of confidence as determined by the L2G scores. L2G scores above 0.1 are shown. IL-1β^high^ genes are shown in red, IL-1β^low^ genes are in blue. **b**) Forest plot showing direction of effect of the rs748670681 variant in the *TNRC18* gene locus associated with immune-mediated diseases in the Finnish population. The minor allele T of rs748670681 concomitantly increases risk for 7 diseases (OR > 1) and decreases risk (OR < 1) for 5 diseases at genome-wide significance (*P* value < 5 × 10^-8^). L2G scores for the *TNRC18* gene in the tile plot in panel A were derived from FinnGen GWAS in data freeze (DF) 6 (n = 260,405), whereas the odds ratios (ORs) in the forest plot in panel B are based on data from DF9 GWAS (n = 377,277), where additional genome-wide significant associations with rs748670681 were detected with increased sample sizes (e.g., multiple sclerosis) that were not detected in DF6. CID, chronic inflammatory diseases; AS, ankylosing spondylitis, CD, Crohn’s disease; PSO, psoriasis; PSC, primary sclerosing cholangitis; SLE, systemic lupus erythematosus; SS, systemic sclerosis; UC, ulcerative colitis.

Some loci, including those containing genes such as *SH2B3*, *IL12RB2*, and *TNFAIP3*, show association with multiple diseases, reflecting possible pleiotropy of these genes that influence disease risk through modulation of IL-1β production. For example, *TNFAIP3* encodes the AP20 enzyme, a known regulator of NFκB-mediated immune activation^36^ and pro-IL-1β processing^37,38^. The association signals at the *TNFAIP3* gene locus have been resolved to *TNFAIP3* using experimental^39–41^ or *in silico*^42^ methods for some of the immune-mediated diseases.

Concurrently, the screening results facilitated prioritization of candidate target genes in unmapped risk loci. For example, *TENT4A* can potentially be prioritized as a candidate gene for multiple sclerosis (MS), given the role of IL-1β in MS^43^. *TENT4A* stabilizes and increases the half-life of mRNA molecules by guanylation of the poly(A) tail^44^. Our screen shows that loss of *TENT4A* results in reduced IL-1β (IL-1β^low^), suggesting that it directly or indirectly contributes to IL-1β production. A common SNP (rs34681760) in the promoter of *TENT4A* is significantly (*P* = 2.0 x 10^-11^) associated with MS risk^45^. This SNP is linked to *TENT4A* with a high L2G score (0.73) and is significantly associated with the expression level of *TENT4A* in monocytes^46^ (*P* = 6 x 10^-30^) and dendritic cells^47^ (*P* = 2 x 10^-15^). However, no experimental evidence currently exists for the biological mechanisms underlying this association signal. These data shed light on potential mechanistic links of such genetic associations and facilitate hypothesis generation for further investigation.

### Genetic pleiotropy of the *TNRC18* gene locus in the Finnish founder population

One of the statistically significant (Q-value < 0.35) genes identified in our CRISPR screen codes for a poorly characterized, intracellular protein TNRC18. The *TNRC18* gene locus has been previously associated with several autoimmune and inflammatory diseases in the Finnish and Estonian populations^32^. The association signal is particularly pronounced for inflammatory bowel disease (IBD) (OR = 2.23, *P* = 1.47 × 10^-117^), followed by ankylosing spondylitis (AS) (OR = 2.33, *P* = 7.6 × 10^-53^). Furthermore, the lead SNP rs748670681 is 114-fold enriched in the Finnish population (MAF = 3.6%) and is rare outside Finland and Estonia. The association signal spans a 3-Mb genomic region that contains 30 protein-coding genes (Fig. 4). Because of the strong genetic association with multiple diseases and the fact that this gene is poorly characterized, we decided to investigate this finding further.

**Figure 4:**
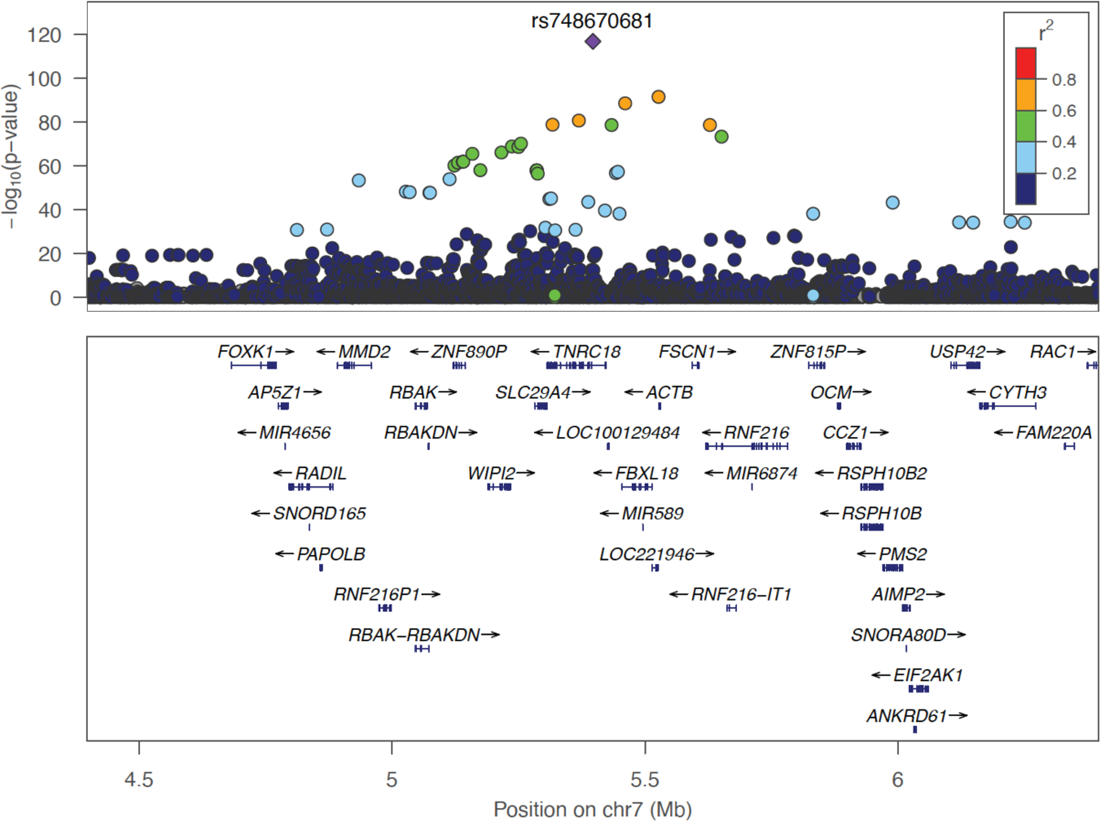
Regional association plot at the *TNRC18* gene locus in IBD. Negative log10 *P* values are shown for variants within a 2Mb region centered at the reference SNP (rs748670681). The reference SNP is marked with a purple diamond, and pairwise LD (*r*^2^) between the reference SNP and other variants are indicated by color. The *r*^2^ values were estimated from high-coverage whole-genome sequences of 8,557 Finns. Both directly genotyped and imputed SNPs are plotted. Genomic coordinates are shown according to the human genome build GRCh38/hg38.

To identify the likely causal variant(s) driving the association signals, we first performed conditional analyses, conditioning on the lead SNP rs748670681 in IBD and type I diabetes, where the variant shows opposite effects^32^. After the conditional analyses rs748670681 remained the sole, disease-associated SNP with genome-wide significance, suggesting that this SNP is driving the association signal for both conditions. To identify other diseases and related phenotypes that might be associated with this SNP, we queried GWAS summary statistics for 2,266 endpoints conducted in 377,277 individuals in the FinnGen study (Supplementary Table 4). The rs748670681 variant showed genome-wide significant (*P* < 5 × 10^-8^) association with ten other immune-mediated diseases. Strikingly, the T allele of rs748670681 showed risk effects for certain inflammatory disorders, whereas it was protective for autoimmune conditions such as Grave’s disease, hypothyroidism, and MS (Fig. 3b).

### The risk allele of rs748670681 alters the expression levels of *TNRC18 and WIPI2*

The immune disease-associated lead variant rs748670681 is located within the second intron of *TNRC18* and overlaps with epigenetic marks for active transcription (H3K27Ac), enhancer activity (H3K4me1), and promoter activity (H3K4me3) in several ENCODE^48^ studied cells including NHLFs, NHEKs, and K562s (Fig. 5a). To test the regulatory activity of this sequence we engineered DNA fragments containing either the risk-associated (i.e., T) or non-risk associated (i.e., C) allele of the rs748670681 variant into a luciferase-based reporter vector. We independently transfected these constructs into Jurkat cells, an immortalized T lymphocyte line, or Caco-2 cells, an immortalized cell line of human colorectal adenocarcinoma cells. The vector harboring the non-risk allele C significantly increased luciferase expression levels compared to the empty vector. However, the vector carrying the risk allele T showed significantly reduced activity compared to the non-risk allele C in both cell types (Fig. 5b and c). This confirms the functional activity of the rs748670681 variant and demonstrates that the risk allele T has decreased transcriptional activity.

**Figure 5:**
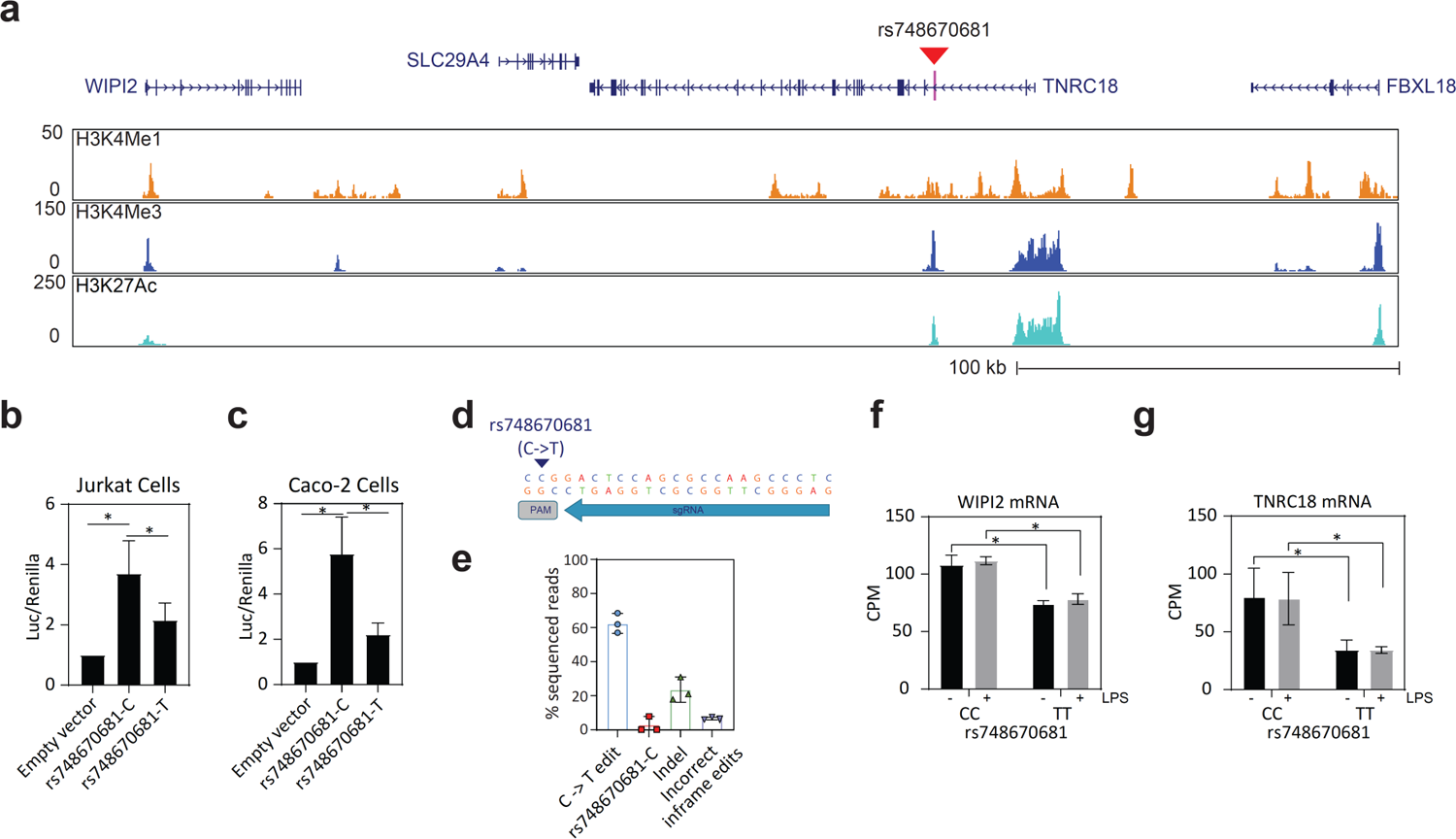
The rs748670681 variant is a transcriptional regulator of the *TNRC18* and *WIPI2* genes. **a**) UCSC genome browser view of the *TNRC18* gene locus. Tracks from top to bottom show RefSeq protein coding genes, H3K4Me1, H3K4Me3, and H3K27Ac marks in NHEKs cells from the ENCODE Project. The rs748670681 variant located inside intron 2 of *TNRC18* is marked with a red triangle. **b-c**) The minor, risk-associated T allele of rs748670681 diminishes luciferase reporter gene expression in Jurkat (**b**) and Caco-2 (**c**) cells compared to the common, non-risk allele C. **d**) Target sequence for HDR-mediated CRISPR-Cas9 editing of the rs748670681 variant sequence. Editing was performed in three independent experiments and quantified using next-generation sequencing with at least 50K sequencing reads. **f-g**) Normalized expression values of *WIPI2* (**f**) and *TNRC18* (**g**) transcripts in cells edited with CRISPR-Cas9 to be homozygous for the T allele of rs748670681. Each bar in graphs represents mean of replicates ± SD (n = 3). Statistical significance was determined using two-tailed, unpaired Student’s *t* test (\**P* < 0.05). CPM, counts per million; Luc, luciferase; PAM, protospacer adjacent motif; LPS, lipopolysaccharide. Statistical significance was determined using two-tailed, unpaired Student’s *t* test (\**P* < 0.05).

To gain a better understanding of the functional effect of the risk allele of rs748670681, we utilized CRISPR-Cas9 mediated homology-directed repair (HDR) and engineered U937 cells to harbor homozygous risk (TT) and non-risk (CC) genotypes. We strategically designed conversion of the C to T (non-risk to risk allele) in our sgRNA (GAGGGCTTGGCGCTGGAGTC) to disrupt the PAM (NGG) sequence and prevent successive Cas9 cutting of our correctly edited genomic sequence. After allowing the cells to recover for 72 hours, we performed next-generation sequencing analysis of the bulk edited cell populations and identified ∼60% genome editing efficiencies across four independent samples (Fig. 5e). We successfully isolated several homozygous clones for the non-risk (CC) and the risk (TT) genotypes, which were subjected to global transcriptomic analysis under PMA differentiated -/+ LPS treatments.

The mRNA expression levels for both *WIPI2* (vehicle: log_2_ fold-change [LFC] = −0.55, *P* < 1.1 × 10^-15^, LPS: LFC = −0.5, *P* < 1.4 × 10^-11^) and *TNRC18* (vehicle: LFC = −0.76, *P* < 0.001, LPS: LCF = −0.87, *P* < 0.001) were reduced in cells engineered to carry the homozygous rs748670681 TT risk genotype compared to the non-risk CC genotype (Fig. 5f and g). The expression of two nearby genes, *SLC29A4* and *FBXL18*, were below reliable detection by RNA-seq in U937 cells. Therefore, the genetic variant rs748670681 appears to exhibit pleiotropic regulatory effects for two genes in the *TNRC18* locus: *TNRC18* and *WIPI2*.

### TNRC18 regulates expression and secretion of proinflammatory cytokines

The identification of *TNRC18* as a genome-wide CRISPR screening hit of IL-1β expression and an abundance of genetic evidence linking it to numerous immune-mediated diseases prompted us to explore its role in myeloid cells in more detail. *TNRC18* is expressed as 10 unique splice isoforms, with the longest spanning 30 exons (2968 AAs) and the shortest 3 exons (99 AAs), each with cell-type specific expression (Supplementary Fig. 1). Although the exact function of TNRC18 is currently unknown, it has been reported to interact with several proteins involved in chromatin remodeling^49–51^, suggesting a potential role for TNRC18 in transcriptional regulation. Within our CRISPR screen, three *TNRC18* guides were enriched in the IL-1β^low^ populations compared to either the presort samples or high IL-1β sorted populations (IL-1β^high^) (Supplementary Fig. 2a). We generated *TNRC18* knockout U937 cells using CRISPR-Cas9 with Inference of CRISPR Edits (ICE) score of 80% (Supplementary Fig. 2b) and confirmed the LPS-dependent reduction of IL-1β in *TNRC18* knockout cells relative to non-edited cells by flow cytometry (Supplementary Fig. 2c and d). *TNRC18* was also required for maximal secretion of proinflammatory cytokines IL-1β, IL-6 and TNF-α in LPS-treated U937 cells (Supplementary Fig. 2e-g). Finally, Cas9 ribonucleoproteins (RNPs) targeting *TNRC18* in primary macrophages from human donors resulted in reduced levels of secreted IL-1β, IL-6, and TNF-α in response to a range of LPS doses (Supplementary Fig. 3). Collectively, these results demonstrate the requirement for *TNRC18* in LPS-dependent activation and secretion of proinflammatory cytokines in immortalized and primary human myeloid cells.

### The global role of rs74867068 in proinflammatory gene programs in myeloid cells

To assess whether U937 cells carrying the homozygous TT risk genotype of rs74867068 demonstrated diminished regulation of proinflammatory gene expression we generated global transcriptomic profiles using RNA-seq. We compared the expression of individual genes between cells homozygous for the CC non-risk genotype versus those engineered to carry the homozygous TT risk genotype with and without LPS stimulation. Compared to cells homozygous for the CC genotype, TT cells treated with LPS showed overexpression of 658 genes with LFC > 1 and adjusted *P* value < 0.05. Conversely, 433 genes were downregulated at the same threshold (Fig. 6a). Genes elevated in cells homozygous for the TT risk genotype are enriched for interferon signaling pathways and defense to viral responses, including *IFIT3, OAS3*, *IFI6* (Fig 6b.), while genes exhibiting decreased expression include *IL1B*, *CCL4*, *CXCL8* and genes participating in pathways responsible for regulating immune cell responses, cell migration, and proliferation (Fig 6c). Selected genes among the up- and down-regulated genes are shown in Fig. 6d-e and highlight increases in interferon-responsive genes (*IFIT3*, *OAS3*, and *IFI6*) and decreases in inflammatory cytokines (*IL1B*, *CCL20*, and *CCL4*). These findings suggest that the rs74867068 allele affects the expression of *TNRC18*, *WIPI2* and consequently many proinflammatory cytokines and interferon responsive genes (Fig. 6f).

**Figure 6:**
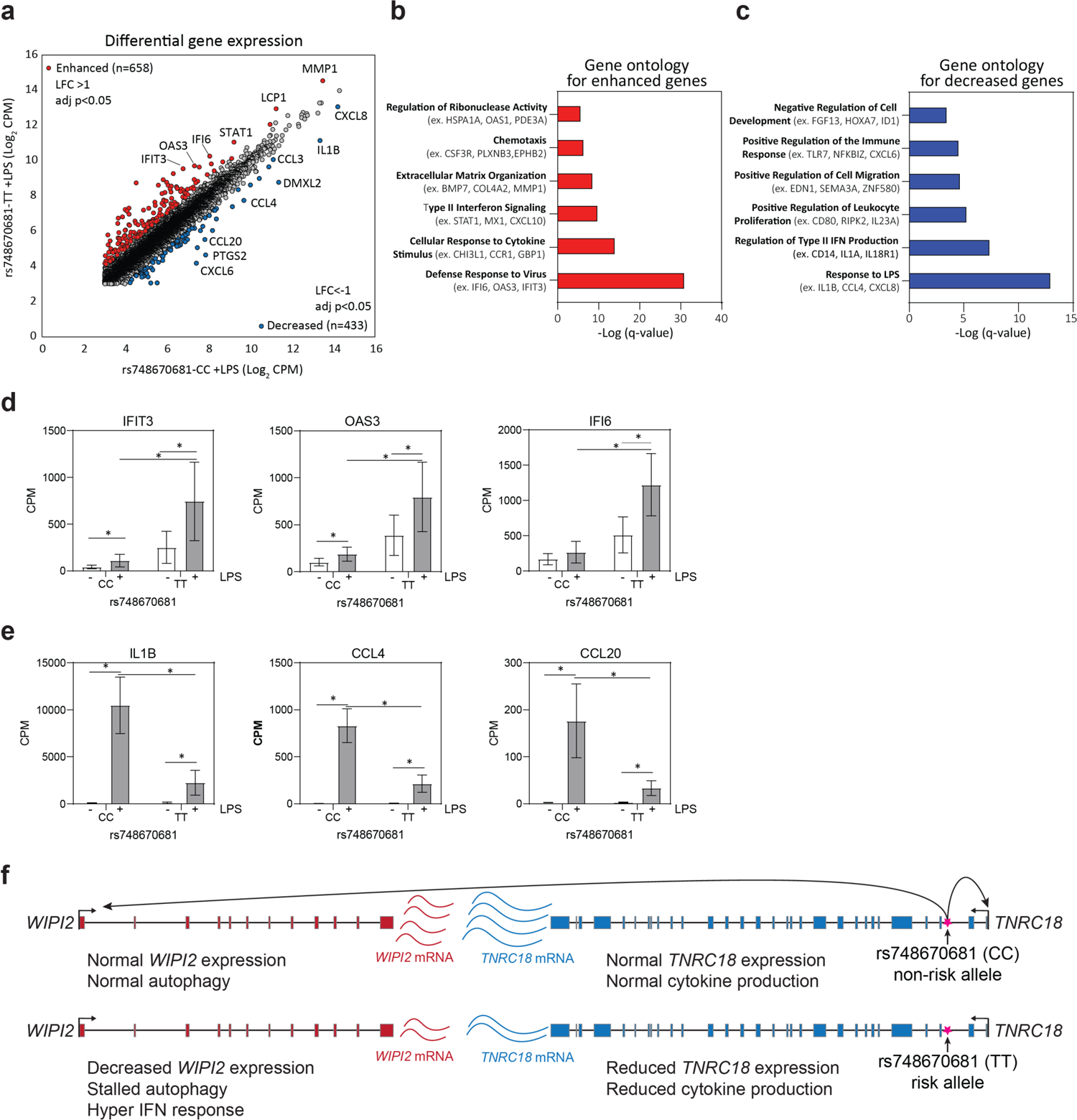
The risk allele T of rs748670681 alters *TNRC18* and global expression signatures. **a**) Differentially expressed genes in U937 cells edited with CRISPR-Cas9 to be homozygous for the T allele of rs748670681 and treated with LPS compared with LPS-treated rs748670681-CC cells. **b-c**) Pathway enrichment using gene ontology terms for biological processes. Selected pathways are shown for enhanced (**b**) and decreased (**c**) genes ranked by -log(q-value). **d-e**) Risk allele T of rs748670681 increases expression of the *IFIT3*, *OAS3*, and *IFIH6* genes (**d**), but decreases expression of *IL1B*, *CCL4* and *CCL20* genes (**e**). Effect of rs748670681 on gene expression was assessed with and without LPS stimulation. (**f**) Schematic of rs748670681 regulation of *WIPI2* and *TNRC18*. LFC, log_2_(fold-change); CPM, count per million; IFN, interferon. Statistical significance was determined using two-tailed, unpaired Student’s *t* test (\**P* < 0.05).

### *TNRC18* and *WIPI2* regulate distinct sets of genes that overlap with genes altered by rs74867068

The observation that the rs74867068 variant modulates the expression levels of both the *TNRC18* and *WIPI2* mRNAs prompted us to examine whether we could assign the gene regulation patterns defined in U937 cells carrying the TT genotype of rs74867068 to knockout of either *TNRC18* or *WIPI2*. Knockout of *WIPI2* leads to the increased expression of 684 genes with the overlap of 241 genes with rs74867068-TT U937 cells TT (*P* = 5.77 × 10^-81^) (Fig. 7a) that are highly enriched for interferon response genes predicted to be regulated by transcription factors STAT1 and IRF1 (Fig. 7c and e) and include *IFIT3*, *OAS3*, and *IFI6* (Fig. 7g). In contrast, knockout of *TNRC18* leads to overlap of 48 down-regulated genes with rs74867068-TT U937 cells (Fig. 7b) (*P* = 0.0003) that are highly enriched for cytokine production predicted to be regulated by the transcription factor NFκB (Fig. 7d and f) and exemplified by the proinflammatory cytokine genes *IL1B*, *CCL4*, and *CCL20* (Fig 7h). These results demonstrate that knockout of *WIPI2* leads to enhanced expression of interferon-responsive genes, while TNRC18 controls the expression of proinflammatory cytokines.

**Figure 7:**
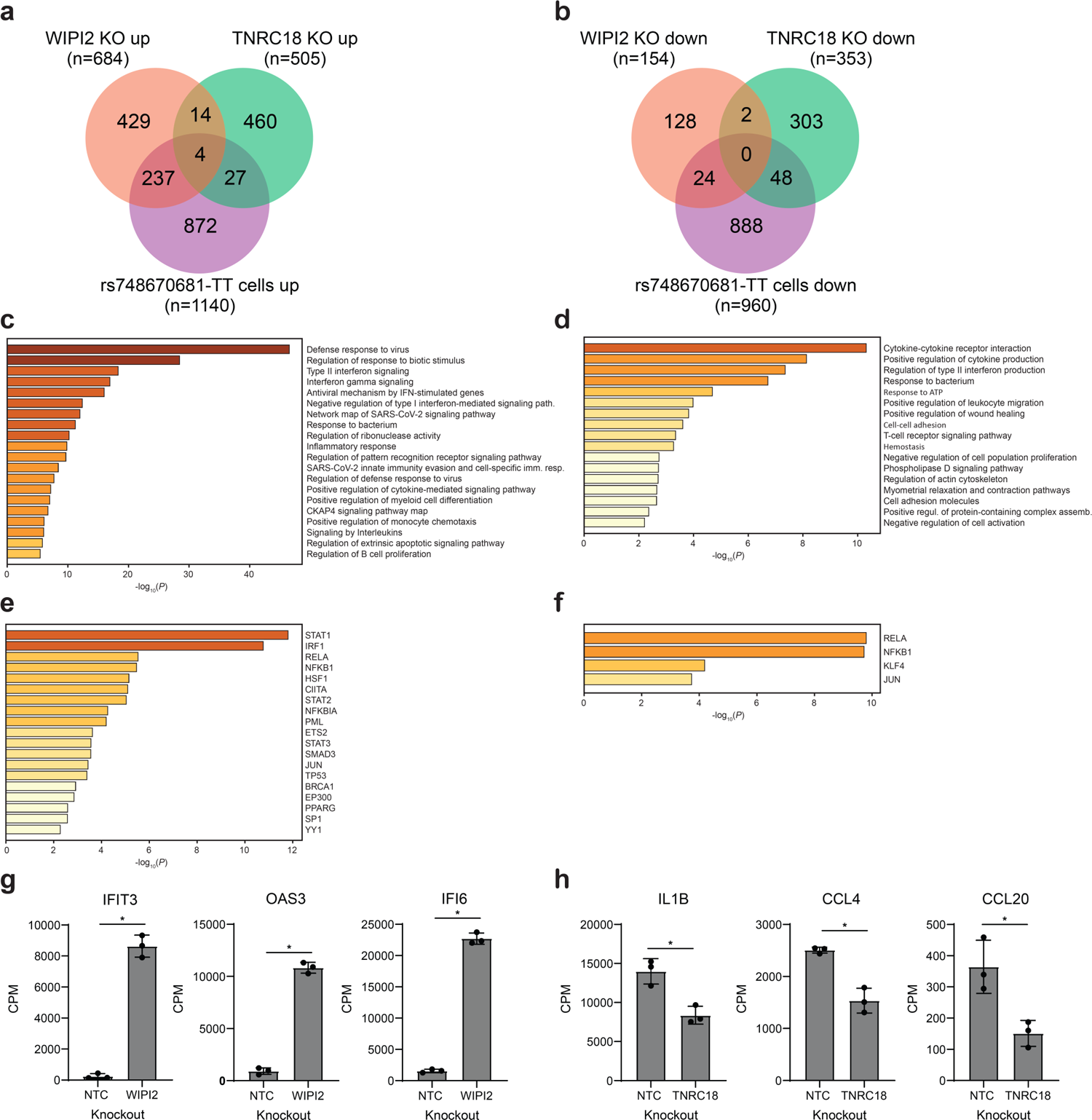
Genes and pathways affected by loss of *WIPI2* and *TNRC18* genes in U937 cells. **a-b**) Number of overlapping genes up-regulated (**a**) and down-regulated (**b**) in *WIPI2* KO, *TNRC18* KO cells, and cells homozygous for the T allele of rs748670681 (LFC > 0.58, adjusted *P*-value < 0.05). **c-d**) Biological pathways enriched in the overlap between genes up in both *WIPI2* KO and rs748670681-TT cells (n=241) (**c**) and enriched in the overlap between genes down in both *TNRC18* KO and rs748670681-TT cells (n=61) (**d**). Motif enrichment in the overlap between genes up in both *WIPI2* KO and rs748670681-TT cells (n=241) (**e**) and enriched in the overlap between genes down in both *TNRC18* KO and rs748670681-TT cells (n=48) (**f**). Selected genes which are increased when *WIPI2* is knocked out (**g**) and decreased when *TNRC18* is knocked out (**h**). Each bar in graphs represents mean of replicates ± SD (n = 3). Statistical significance was determined using two-tailed, unpaired Student’s *t* test (\**P* < 0.05). KO, knock-out; CPM, counts per million; NTC, non-template control.

## DISCUSSION

Therapeutic targets with causal relationships to human diseases demonstrate increased success rate in clinical development. Perhaps the clearest proof of causality for any non-communicable human disease comes from genetics. Since 2005, thousands of GWAS have uncovered genetic variation influencing risk for human complex diseases, including many with alterations in the immune system^52,53^. Most GWAS disease-associated SNPs localize to noncoding regions, suggesting that they affect gene function through regulation of gene expression. Following GWAS, various computational and experimental approaches need to be undertaken to connect statistically significant signals in non-coding regions to causal variants and genes, such as colocalization with expression quantitative trait loci (eQTL), chromatin accessibility or interaction analysis. However, the molecular and cellular mechanisms by which the causal genes contribute to disease risk do not always become evident, especially for those with no known biological function. Linkage disequilibrium can further complicate the identification of true causal alleles, since it is not uncommon to observe eQTL SNPs affecting more than one gene. Unbiased CRISPR loss-of-function screens in disease-relevant cells and states offer a complementary and powerful approach to identify disease-associated genes under GWAS LD peaks, simultaneously providing mechanistic insights into pathogenic processes. The IL-1β cytokine and macrophages are among the most critical drivers of inflammation underlying immune diseases. Here we describe the first genome-wide CRISPR knockout screen followed by arrayed validation to identify regulators of intracellular IL-1β content in the human monocytic cell line U937 under stimulation conditions. Our screen identified many known modulators of IL-1β production including the *IL1B* gene itself. The screen also identified novel genes, providing a list of potential target candidates for further investigation. Most importantly, 57 (19%) of 295 screening hits reside in genomic loci that have been associated with various inflammatory or autoimmune diseases.

We further characterized one of the most significant GWAS risk loci in the Finnish population. The 7p22.1 chromosomal locus was previously found to modify the risk for several autoimmune diseases in the Finnish and Estonian population^32^. The derived allele of the noncoding SNP rs748670681 concomitantly increases the risk for certain diseases and decreases the risk for others. Statistical fine mapping of the risk haplotype suggested that the intronic lead SNP rs748670681 is causal and the *TNRC18* gene, which harbors this SNP, was identified as a significant hit in our CRISPR knockout screen, suggesting that *TNRC18* could be the effector gene of this GWAS signal. To elucidate the functional role of this variant and to identify the causal genes at the locus, we generated a CRISPR-Cas9 edited cell model carrying the homozygous risk genotype and showed the effect of the risk genotype on the expression of *TNRC18* and, also surprisingly on the adjacent *WIPI2* gene.

The precise function of the TNRC18 protein remains uncharacterized. In high-throughput proteomics assays, TNRC18 was found to interact with several other proteins including regulators of chromatin structure (HDAC1, FAM60A, SIN3A, SIN3B, SAP30, PRDM5, HIST1H1E)^49–51,54–58^ and transcription factors (AR, ESR2, SOX2)^59–61^. Moreover, TNRC18 contains a bromo-adjacent homology (BAH) domain found in a variety of proteins playing roles in chromatin remodeling and transcriptional silencing^62^, as well as a Tudor domain that recognizes and binds to methylated lysine and arginine residues allowing them to function as readers of the epigenetic landscape^63^. Furthermore, *TNRC18* was identified as a top positive hit that confers resistance against mycobacterium infection in CRISPR knockout and knockdown screens in human monocytic THP-1 cells (Lai et al.)^64^. Interestingly, some of the same pathways (e.g., type I interferon signaling) and top gene hits (*JAK1*, *SHOC2, JUNB, AHR, ARNT*) were common between our and the Lai et al. study. Consistent with the predicted function, our findings confirm gene regulatory activities of the TNRC18 protein and support a role in the regulation of inflammatory signaling, potentially explaining genetic associations with multiple immune diseases.

*WIPI2* codes for a component of the autophagy pathway^65^. Genetic variation in autophagy genes and perturbation of the autophagy pathway, in general, have been implicated in susceptibility to immune diseases^66,67^. The risk allele of rs748670681 diminishes cytokine expression, yet concurrently enhancing expression of interferon signaling pathway genes. The latter finding is noteworthy, since the risk allele demonstrates protective effects for multiple sclerosis, whose treatment has been interferon injections^68^. In validation studies, reduced expression of cytokine genes and induction of the interferon signaling genes appear to result from loss of *TNRC18* and *WIPI2*, respectively. The risk allele decreases the expression of both genes. The opposing effects of the rs748670681 variant on disease risk may be attributed to combined perturbations of both *WIPI2* and *TNRC18* expression or due to their separate effects.

In conclusion, our study defines global regulation strategies for IL-1β production in macrophage cells and identifies *TNRC18* as a novel modifier of LPS-dependent transcription. We provide evidence for a causal effect of the rs748670681 variant on chromosome 7p22.1 in various immune diseases by modulating *TNRC18* and *WIPI2* gene expression. Further studies are necessary to gain a deeper understanding of the pleiotropic effect of the rs748670681 variant, which contributes to increased risk for some immune diseases, simultaneously lowering the risk for others.

## METHODS

### Cell culture

U937 cells were maintained in RPMI 1640 (Gibco) supplemented with 10% heat inactivated fetal bovine serum (FBS, Gibco) and 1% Pen/Strep (Gibco). Cells were differentiated by addition of 10 ng/ml phorbol 12-myristate 13-acetate (Sigma, P1585) for 24 hours and treated with 100 mg/ml lipopolysaccharide (LPS) (Sigma, L26375MG) or Nigercin (Invivogen, tlrl-nig-5) for designated times. Primary human CD14+ monocytes (BioIVT) were maintained in RPMI 1640 (Gibco) with sodium pyruvate (ThermoFisher, 11360070), non-essential amino acids, HEPES (ThermoFisher, 15630080), 1% Pen/Strep (ThermoFisher, 15070063), beta-mercaptoethanol (Sigma, M6250), GlutaMAX (ThermoFisher, 35050079), 10% FBS (Gibco), and 100 ng/mL recombinant human M-CSF (Peprotech, 3002510UG) at 37°C, 5% CO_2_. Jurkat cells were grown in RPMI 1640 medium with 10% heat-inactivated FBS, and Caco-2 cells were cultured in DMEM supplemented with 10% heat-inactivated FBS (all from Gibco, Life Technologies) in a 37°C humidified 5% CO_2_ incubator.

### Genome-wide CRISPR screening

120 million U937-LC3-Cas9 cells, U937 cells that stably express Cas9 and a LC3 autophagy reporter, were infected with the Brunello sgRNA library at multiplicity of infection (MOI) = 0.3 in three biological replicates. The Brunello CRISPR-Cas9 knockout library contains 76,441 sgRNAs targeting 19,114 genes with 1,000 non-targeting controls^34^. Following puromycin selection, cells were differentiated with PMA and treated with LPS for 24 hours. 40 million cells were collected with trypsin as presort for genomic DNA extraction for each replicate. Greater than 120 million cells per replicate were then processed and stained for intracellular IL-1β at 10 million cells/ml using the protocol mentioned in the Flow Cytometry section. Cell populations expressing the top and bottom 10% IL-1β levels were enriched using a FACSAria Fusion cell sorter (BD Biosciences). At least four million IL-1β^low^ and IL-1β^high^ cells were collected for genomic DNA extraction using Quick-DNA FFPE Kit (Zymo Research). PCR reactions containing up to 10 μg genomic DNA in each reaction were performed using ExTaq DNA Polymerase with primers to amplify the sgRNAs. Samples were then purified with SPRIselect beads, mixed and sequenced on a NextSeq 500 by 75-bp single-end sequencing.

### Screen analysis

Sequencing data were demultiplexed and reads with sgRNA sequences were quantified using custom Perl scripts. Given that the position of the sgRNA could vary per read, the primer sequences flanking the sgRNA were considered. Sequences in between the flanking primers were extracted and then compared to sequences in the sgRNA library. Only sequences with no mismatches were used in the calculation of guide-level read counts. Samples with a minimum of 80% reads mapping to a sgRNAs were considered for further analysis. Through the *edgeR* Bioconductor package^69^, reads were normalized between samples using the trimmed mean of M values (TMM) method^70^. Differential analysis of guide level counts was performed using the *limma* R Bioconductor package^71^. Gene level effects were considered as the median log fold change of the four guides associated with each gene. Statistically significant genes were identified by aggregating differential sgRNAs ranked consistently higher at gene level using the robust rank aggregation (RRA) method^72^.

### Flow cytometry

Cells were harvested and washed twice with FACS buffer (PBS + 1% FBS), incubated with IC Fix buffer (eBioscience, Cat #88-8824-00) in dark for 30 minutes at room temperature, washed twice with FACS buffer, resuspended in 1X Permeabilization buffer (Ebioscience, Cat #88-8824-00) for 10 minutes and washed twice with 1X perm buffer. Cells were then resuspended in 100 mL/well of FC-block (Biolegend, 422302) (1:100 in Perm buffer) for 15 minutes at room temperature in the dark. Without washing, directly add 100 mL/well of IL-1β (Biolegend, 508208) at 2X (1:50 in perm buffer) to the FC-block currently in well (1:100 final dilution) and incubate at RT for 15 minutes in the dark. Cells were then washed twice in 1X Permeabilization Buffer, resuspended in 200 ml of FACS buffer and fluorescence was measured using a BD LSRFortessa Cell Analyzer. Data was analyzed using the Flow-Jo (v.10) analysis software.

### Arrayed screening and cytokine detection

Library of arrayed guide lentivirus was obtained from Broad Institute targeting screening hits with 2-3 guides/gene. U937-Cas9 clonal cell line (U937-AA7F9) were seeded at 0.05M cells/100 μl/well in a 96-well plate in presence of 8 μg/ml polybrene and 10 μg/ml blasticidin using Multidrop Combi. The cells were then transduced with guide lentivirus using Bravo automated liquid handler from Agilent in three different replicates and incubated at 37°C. Two wells of each screening microplate were transduced with positive and negative control guide lentivirus. After overnight incubation transduction media was changed into selection media containing 10 ug/ml blasticidin and 2 μg/ml puromycin and returned to 37°C incubation. After 3 days of puromycin selection, lentivirus transduced cells were thoroughly mixed and equally divided into 4 different 96 well plates – to be used downstream for IL-1β flow assay and cytokine secretion assay run in duplicate. For IL-1β flow assay, cells were further treated with 100 ng/ml PMA for 24hrs, rested for 24hrs in PMA-free media followed by 24hr treatment with 100 ng/ml LPS, after which cells were harvested for IL-1β flow assay. For cytokine secretion assays, cells were treated with 25 ng/ml PMA for 3 days, rested in PMA-free media for 4 days and then treated with 1.5 mg/ml LPS with and without 1 μM Nigericin for 24hrs. After LPS treatment cell supernatants were collected and analyzed for IL-1β secretion by MSD assay. Puromycin selection was continued throughout PMA and LPS treatments and all media changes and compound treatments were performed using a Bravo and Multidrop Combi respectively. The sgRNA sequences are provided in Supplementary Table 8.

### Cytokine secretion assays

Cells were treated as previously described and supernatants were collected from three independent samples. Cytokines were assessed using the Proinflammatory Panel 1 (human) kit (Meso Scale Diagnostics, K15049G) according to the manufacturer’s instructions. Cytokine levels were measured using the MESO QuickPlex SQ120 and concentrations were calculated using a standard calibration curve.

### Electroporation and dual luciferase reporter assay

gBlocks with selected SNP-containing fragments (IDT) were cloned into pGL4.10 plasmid (Promega). For the luciferase reporter assay, 150K Caco2, Jurkat cells were nucleofected with 50 ng of renilla vector along with 170 ng of luciferase vectors using the Lonza Nucleofector according to the manufacturer’s protocol. Twenty-four hours after transfection, cells were washed once with cold 1× PBS and the luciferase activities were measured with Perkin Elmer Envision using Promega Dual Luciferase Assay kit (Promega, E1960). The firefly luciferase activity was normalized to renilla luciferase activity for each well. All the luciferase activity measurements were performed in 7 replicates for each condition. The Student’s *t*-test was applied to estimate the statistical significance of the difference in luciferase activities between the two alleles.

### Primary macrophage CRISPR-Cas9 mediated genome editing

CD14+ monocyte cells were thawed and plated in growth media. On day 5, adherent cells were collected over ice using a cell lifter. The cells were washed once with PBS and resuspended in P3 Primary Cell solution (Lonza). RNP complexes were formed by mixing Cas9 protein (IDT, 1081058) with the indicated sgRNA and incubating at room temperature for 10 minutes. RNP complexes were then added to the adherent cell solution and electroporated using the Lonza 4D-Nucleofector and protocol CA-137. The cells were then transferred into media and incubated at 37°C, 5% CO_2_. After 3 days, cells were treated with LPS for times indicated.

### RNA isolation and qPCR

Total RNA was isolated using the Direct-zol RNA Miniprep Plus kit (R2072, Zymo Research) followed by cDNA synthesis with the SuperScript III First Strand Synthesis kit (11752-050, Thermo Fisher) according to the manufacturer’s instructions. qPCR was performed with PowerTrack Sybr Green Master Mix (A46109, Applied BioSystems) with the following cycling conditions: 95°C for 2 min then 40 cycles of 95°C for 5sec and 68°C for 30 sec. A melting curve was conducted to verify the absence of non-specific amplification. The Ct values were normalized to *RPLP0* to control for transcript levels. The primer sequences are provided in Supplementary Table 9.

### RNA sequencing

500 nanograms of total RNA were used for NEBNext® Ultra II Directional RNA Library Prep Kit (NEB # E7760L). Poly-A selection (NEBNext Poly (A) Magnetic Isolation Module; NEB # E7490) and cDNA synthesis were performed according to NEB protocol. The NEBNext UMI RNA Adaptors were diluted 50-fold in ice-cold UMI Adaptor dilution buffer. Ligation reaction was purified using Ampure XP beads. (Beckman Coulter, # A63881) and the adaptor ligated cDNA was amplified with 12 cycles of PCR with NEBNext Primer Mix (96 Unique Dual Index UMI Adaptors RNA set1, (NEB #E7416). Equimolar amounts of the samples were then pooled and sequenced using NextSeq 500/550 70 base read lengths in paired-end mode.

### RNA-Seq data analysis

An average of 23.7 million paired-end reads were generated. FastQ files were aligned to human genome reference using STAR^73^ (v2.7.10b) with GRCh38 primary assembly (v2.7.4a) gene annotations. Gene expression levels were counted using featureCounts^74^ (v2.4.3). Differential expression was compared using the test in *limma* R package^71^ (v3.54.2). *P* values were adjusted using the Benjamini and Hochberg method. The significance of the overlap of differentially expressed genes was calculated using the hypergeometric distribution with the phyper function in the R statistical package (v4.2.2). Gene enrichment and pathway analyses were performed using Metascape^75^ (http://metascape.org).

### Design and synthesis of gRNA and HDR repair template

To generate U937 cells carrying the *TNRC18* risk allele, we designed CRISPR/Cas9 gRNA targeting upstream of the mutated allele in IBD on the reverse complementary strand so that the middle G of the PAM corresponds to the C to be edited in the leading strand (Fig. 5d). This prevented re-editing by Cas9 after the initial repair because the PAM was destroyed. The homology-directed repair template contained a T instead of a C franked by the DNA sequence around the allele to be edited. The gRNA was synthesized as a 100-nucleotide containing the *tracrRNA* and 20 nucleotide spacer RNA (IDT). The HDR repair template was synthesized as 85 nucleotides single-stranded DNA with phosphorothioate modifications at 5 and 3’ ends to protect it from exonuclease degradation.

### Nucleofection of Cas9/gRNA RNP complex and HDR repair template

RNP complex containing gRNA and Cas9 protein (IDT, Alt-R® S.p. Cas9 Nuclease V3, Cat #: 1081058) were prepared by mixing 150 pmol gRNA and 125 pmol of the Cas9 enzyme in 1.5 µL PBS to a total 5 µL reaction volume per well of a 96 well plate. The reaction was incubated at room temperature for 20 minutes. U937 cells (100,000 cells/well) were washed twice with PBS and resuspended in a solution containing a 4.5:5 mixture of nucleofection buffer (Buffer SE, Lonza Lot #: S-09279) and supplement (Lonza, Lot #: 09588). The Cell suspension was mixed with the 5 µL of RNP complex, 100 µM HDR template, 100 µM Alt-Cas9 electroporation enhancer (IDT cat# 10007805), and Phosphate buffered saline (PBS) to a final volume of 30 µL. The nucleofection mixture was transferred to a 96-well Nucleocuvette (25 µL/well) and electroporated using Lonza 96-well Shuttle following the FP-100 program. After electroporation, the cells were transferred to a 96-well tissue culture plate containing 1.0 µM Alt-R HDR Enhancer V2 (IDT, Cat #: 10007921) in 200 µL of cell culture media. The cells were incubated at 37°C in a tissue culture incubator for 24 hours. After 24 hours, the media was changed to regular U937 culture media. Cell culture media was changed every 2-3 days until the cells were confluent.

### Characterization of isogeneic cell lines

Genomic DNA was isolated using the Quick DNA Miniprep kit (D3025, Zymo Research) according to the manufacturer’s instructions. PCR was performed with the Phusion High-Fidelity PCR kit (E0553L, New England BioLabs) with the following cycling conditions: 98°C for 3min then 35 cycles of 98°C for 45 sec, 71°C for 30 sec, 72°C for 90 sec, then concluding with 72°C for 10 min. The PCR primer sequences are TNRC18 For1: GATCCCAGTCCAGCCTCTGA and TNRC18 Rev1: GAGGCACTGAGAACGGCACT. PCR product was purified using AMPureXP beads (A63880, Beckman). Nested PCR was performed on the purified PCR amplicon with the following cycling conditions: 98°C for 3 min then 35 cycles of 98°C for 30 sec, 69°C for 30 sec, 72°C for 60 sec, then 72°C for 10 min. The nested PCR primer sequences are: TNRC18 For3: CTTTGGGGTACATGTGGGGA, and TNRC18 Rev3: TTGGGCCGCATGGATGAAC. Amplicons were purified by AMPureXP beads (A63880, Beckman) and submitted to Azenta Life Science for Sanger sequencing on individual clones, CRISPR-Cas9 nucleofected samples, or NGS on bulk edited populations. A successful biallelic edit was indicated by a clean T instead of a C at the specified locus, as visualized by Geneious Prime software. Three different clones were used in the subsequent experiments.

### Open Targets L2G scores

The Open Targets project has performed systematic fine-mapping and gene prioritization at over 130,000 published GWAS loci from the GWAS catalog, the FinnGen study and the UK Biobank^35^. The L2G algorithm trained on a set of unequivocally mapped causative genes using a gradient boosting model has enabled the ranking of genes within each GWAS locus by integrating statistical fine-mapping and various epigenetic signatures as predictive features.

Multiple tables that contain these data from the October 2022 (22.10) release were downloaded in bulk from Google BigQuery^76^. We wrangled the data by merging various tables and harmonizing the phenotype descriptions across different GWAS (Supplementary Table 3). We then queried the L2G analysis output for the top 295 genes identified in our genome-wide screen CRIPSR. Where more than one study linked the same gene to the GWAS signals, we selected the highest L2G score among those studies for plotting. Top L2G scores between genes and phenotypes above 0.1 were plotted using the *ggplot2* R package (v3.3.6).

### FinnGen GWAS

FinnGen is a large public-private partnership aiming to collect and analyze genome and health data from 500,000 Finnish biobank participants^32^. Endpoint definitions derived from electronic health records and curated by clinical expert groups are available in https://www.finngen.fi/en/researchers/clinical-endpoints (DF9). Tools, data QC and association analyses are described in-depth in https://finngen.gitbook.io/documentation (DF9). In brief, 20,175,454 imputed variants were tested between cases and controls using the REGENIE^77^ method and software, adjusting for sex, age, 10 principal components and genotyping batch. Summary statistics of GWAS were obtained from the FinnGen Google Cloud Platform. Regional association plot was generated with the stand-alone version of LocusZoom^78^ (v1.3) using the Finnish population-specific LD structure estimated in 8,557 whole-genome sequences of Finns. Forest plot was made using the R package *forestplotter* (v.0.1.7).

### Conditional analysis

Conditional analysis was performed using the COJO approach^79^ (*--cojo-cond* command) implemented in the GCTA software tool (v1.93.2)^80^. For this analysis, we used summary statistics output from the REGENIE GWAS and imputed genotypes from the entire FinnGen cohort for LD estimation within 3 Mb to each direction from the conditioning SNP.

## DATA AVAILABILITY

Summary statistics from FinnGen GWAS used in this study are available for download from https://www.finngen.fi/en/access_results (DF9 - May 11, 2023).

## ACKNOWLEDGEMENTS

We would like to thank the GTECH lab of the AbbVie Genomics Research Center for preparing and sequencing NGS libraries. We want to acknowledge all participants, administration, and investigators of the FinnGen study. We also thank AbbVie employees Nizar Smaoui, Archana Iyer, and Kathleen Smith for critical review of the manuscript.

## AUTHOR CONTRIBUTIONS

F.R. and J.D.S conceived, designed and led the study and drafted the manuscript. S.G., S.P., M.S., E.N., J.T., D.V., S.S., C.A.E., A.K., T.A., performed experiments, V.A., A.M., C.L., J.W., M.M., N.A.M., analyzed the data, K.V.K., A.B., N.C., M.J.F., A.I.D, J.F.W. provided critical feedback. All authors reviewed the manuscript, provided feedback, and approved the manuscript.

