## Supplementary Fig. for "A genome-wide CRISPR screen supported by human genetics identifies the *TNRC18* gene locus as a novel regulator of inflammatory signaling"

### Supplementary Figures

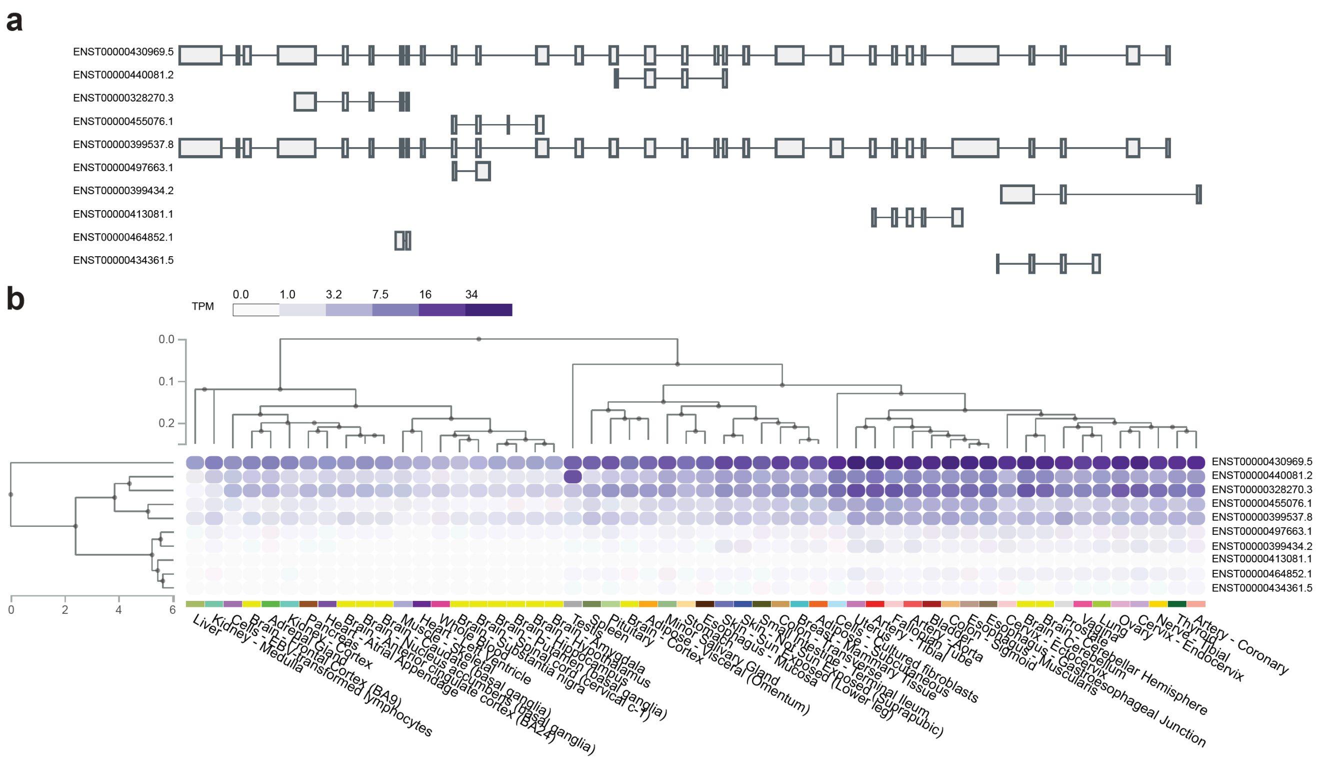


**Supplementary Fig. 1: *TNRC18* gene structure and tissue-specific expression. (from the GTEx database[1])**

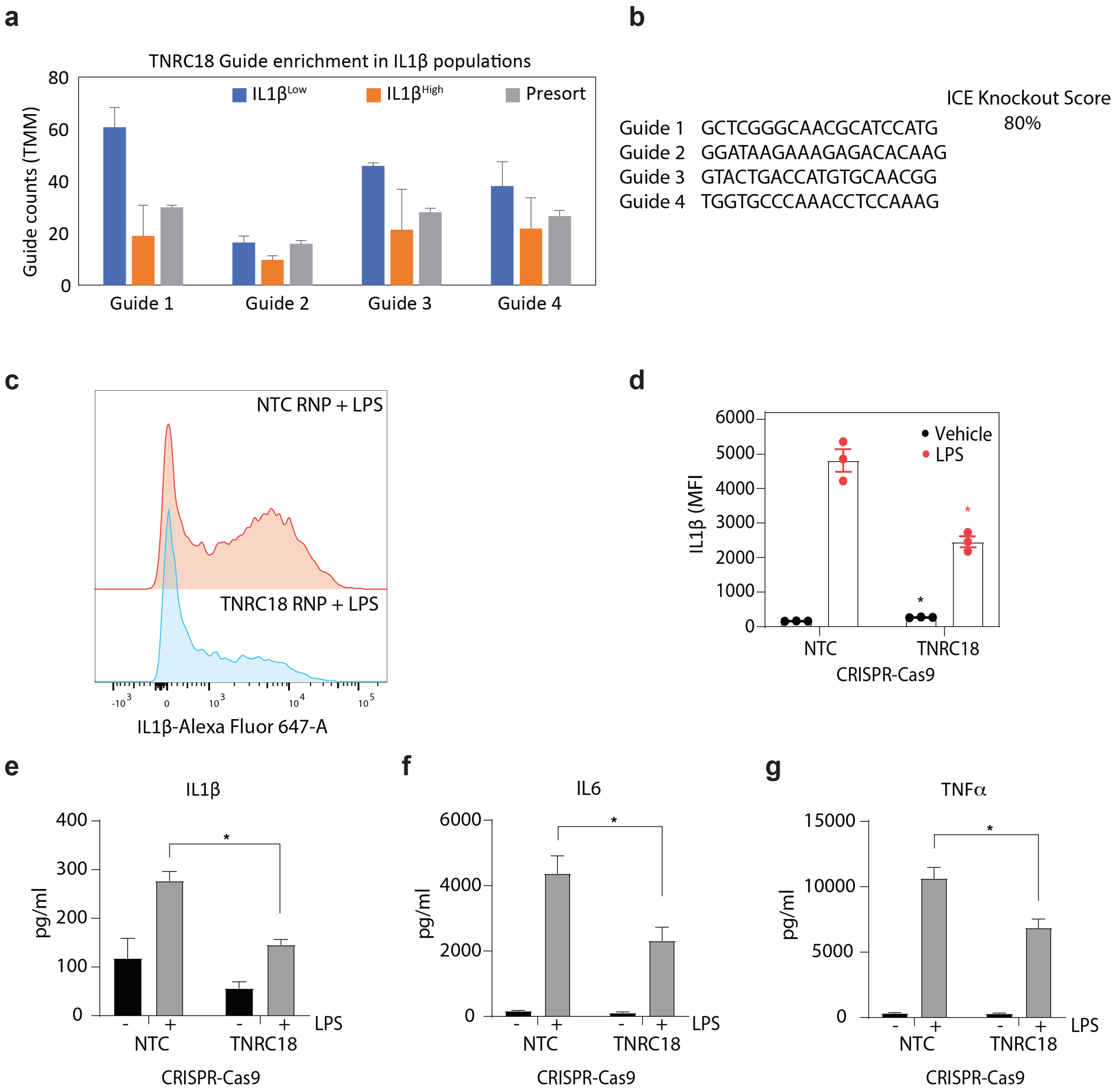


**Supplementary Fig. 2: TNRC18 regulates secretion of inflammatory cytokines.**

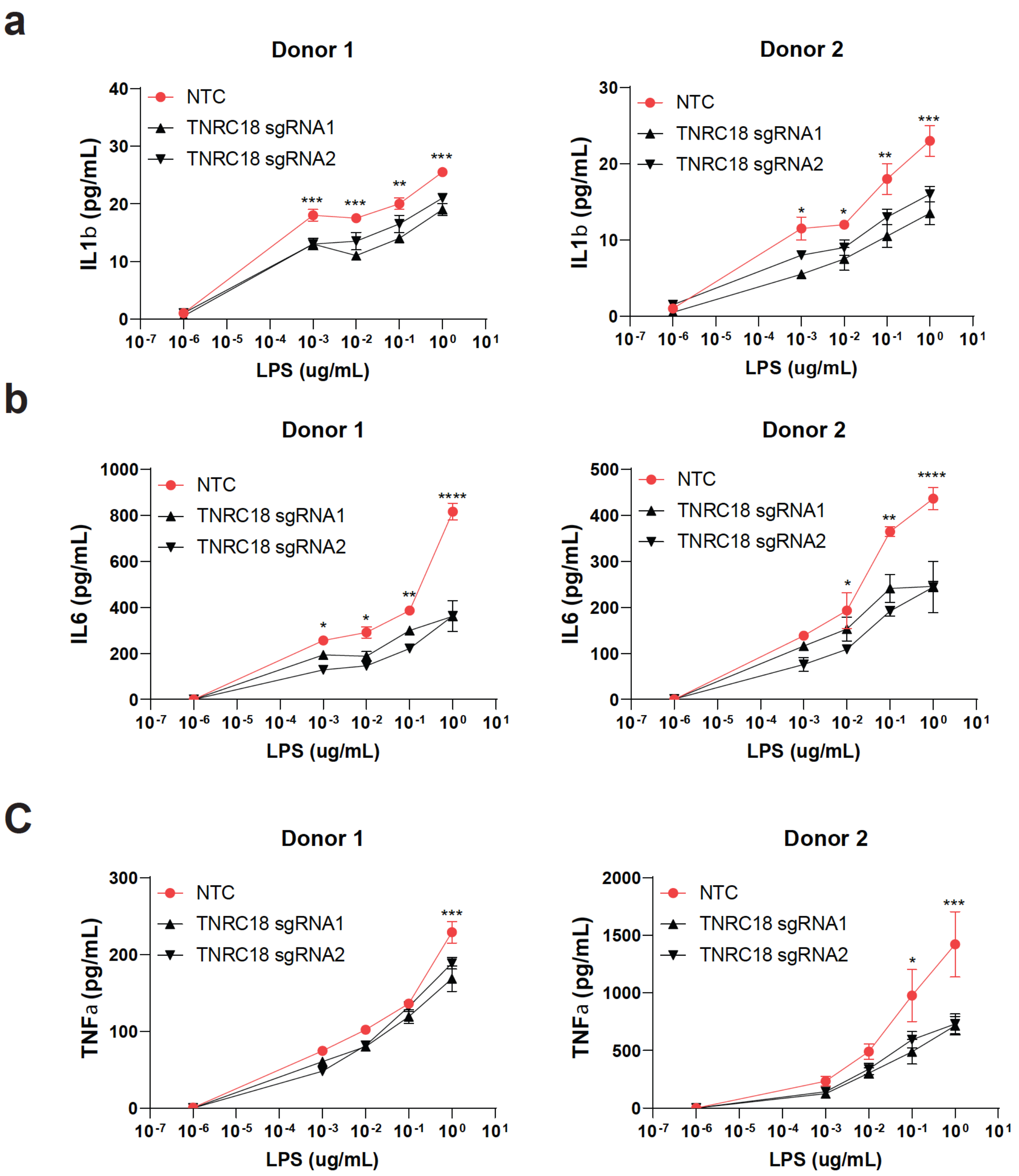


**Supplementary Fig 3. TNRC18 edited CD14+ monocyte derived macrophages produce less LPS induced pro-inflammatory cytokines.**

CD14+ human monocytes from PBMCs were cultured in media containing 100ng/mL M-CSF at 37C and 5% $CO_2$. After five days, cells were harvested and electroporated with indicated RNP (cas9 and sgRNA) and incubated for five additional days. LPS at 100ng/mL was then added to the cells and the supernatant was collected at 24 hours post treatment. Supernatant was assayed for IL1b, IL6 and TNF by MSD.
